# Pdx1 expression in hematopoietic cells activates *Kras*-mutation to drive leukemia in KC (*Pdx1-Cre ; LSL-Kras^G12D/+^*) mice

**DOI:** 10.1101/2022.05.17.492353

**Authors:** Morgan T. Walcheck, Manabu Nukaya, Erik A. Ranheim, Kristina A. Matkowskyj, Sean Ronnekleiv-Kelly

## Abstract

**Background:** *Pdx1* expression in pancreatic lineage cells underlies the utility of the KC mouse model (*Pdx1-Cre; LSL-Kras^G12D/+^*) for understanding how *Kras^G12D^*-mutation drives formation of pancreas cancer precursor lesions and carcinoma. The highly utilized KC model has a reported mortality rate of about 30%, which has been attributed to pancreas cancer, despite lack of substantive evidence. This study describes a novel cause of the early deaths, in which KC mice develop *Kras*-driven T-cell acute lymphoblastic leukemia (T-ALL).

**Methods:** KC mice and control mice underwent histopathologic examination including thymus, liver, spleen, bone marrow and pancreas, and immunohistochemistry (IHC) was used to confirm leukemia development. A reporter strain (Ai14) was used to identify location of *Pdx1-Cre* expression and concomitant mutant-*Kras* activation, which was confirmed using flow cytometry, IHC, immunofluorescence, mRNA analysis, and bone marrow transplant studies.

**Results:** *Pdx1* expression in the hematopoietic compartment of KC mice resulted in Cre-recombinase mediated excision of lox-stop and activation of mutant-*Kras* gene (*Kras^G12D/+^*) in the multipotent progenitor cells (MPP), and subsequent development of *Kras*-mutant T-ALL creating thymic tumors in a subset of mice. Overall, 20% (5/25) of KC mice developed a large thymic tumor due to T-ALL by 9 months of age. Moreover, through isolation and transplantation of pooled bone marrow from KC mice into CD45 congenic mice, 100% of recipient mice were found to develop T-ALL. These results further confirm mutant-*Kras* expression in the hematologic compartment is driving the development of T-ALL in the KC mouse model.

**Conclusions:** These results are an essential consideration for investigators while utilizing this model in pancreas cancer studies, particularly when evaluating factors that may coincidentally enhance the formation of *Kras^G12D^*-driven T-ALL (e.g. transcription factors impacting hematopoietic cells). Finally, the lower penetrance of T-ALL development in KC mice (compared to existing leukemia models) suggest that the KC mouse could be considered as an alternative research model to evaluate onset and factors that exacerbate development of T-ALL.

## BACKGROUND

*Pdx1* expression in pancreatic lineage cells underlies the utility of the KC mouse model (*Pdx1-Cre; LSL-Kras^G12D/+^*) for understanding how *Kras^G12D^*-mutation drives formation of pancreas cancer pre-cursor lesions and ultimately pancreas cancer. This mouse model was first developed in 2003^1^ and has since continued to be highly utilized in pancreas cancer development studies. The KC mouse model is generated by crossing the *LSL-Kras^G12D/+^* (K) mouse to pancreatic and duodenal homeobox 1 (*Pdx1)-Cre* (C) mice resulting in expression of the *Kras*-mutation (*Kras ^G12D^*) in cells expressing *Cre*-recombinase from pancreatic and duodenal homeobox 1 (*Pdx1*) promoter.^1^ *Pdx1* is most highly expressed in pancreas lineage cells, and all KC mice develop pancreatic intraepithelial neoplasia (PanIN)-1 with associated fibro-inflammation by age 5 months. With increasing age, the PanIN lesions progress such that a significant percentage of KC mice develop PanIN-2, PanIN-3 and pancreas cancer by roughly 9 months of age.^1^

Throughout the use of this mouse model, the published survival for KC mice of roughly 70% at 9 months has been attributed to the development of pancreas cancer and consequent lethality.^2^ However, a pancreas-specific cause of lethality can be challenging to identify postmortem due to pancreas enzyme auto-digestion, so diagnosis of pancreas cancer must be confirmed when mice are moribund (i.e. at time of sacrifice as opposed to after mice are already deceased), or there must be evidence of solid organ metastases. While utilizing the KC model, we found that there was an approximate 30% rate of lethality or moribund status requiring euthanasia out to age 9 months, which matches existing literature. However, contrary to prior suppositions, no mice became moribund due to pancreas cancer despite diagnosing pancreas cancer in several of the mice, and there was no evidence for metastatic pancreas cancer in any of the moribund mice. Rather, KC mice became distressed - manifest by respiratory distress or significant weight loss - due to advanced hematologic cancer (T-cell acute lymphoblastic leukemia or T-ALL) resulting in large thymic tumors precipitating respiratory suppression and subsequent death.

This study presents the novel finding that *Pdx1* expression in the lineage negative cells and multipotential progenitor cells (MPP) of the bone marrow (BM) is sufficient to drive Cre expression in this cellular compartment. Thus, in KC mice, *Pdx1* expression results in *Cre-* recombinase mediated excision of lox-stop and activation of mutant-*Kras* gene (*Kras^G12D/+^*) in the hematopoietic cells, and subsequent development of *Kras*-mutant T-ALL with concomitant thymic tumor in a subset of mice. The use of a reporter mouse line and bone marrow transplant study confirmed these results. These results are an important contribution and a notable consideration for investigators utilizing this model in pancreas cancer studies, particularly when evaluating factors that may coincidentally enhance the formation of *Kras^G12D^*-driven T-ALL (e.g. transcription factors impacting hematopoietic cells).

## MATERIALS AND METHODS

### Animals

All animal studies were conducted according to an approved protocol (M005959) by the University of Wisconsin School of Medicine and Public Health (UW SMPH) Institutional Animal Care and Use Committee (IACUC). Mice were housed in an Assessment and Accreditation of Laboratory Animal Care (AALAC) accredited selective pathogen-free facility (UW Medical Sciences Center) on corncob bedding with chow diet (Mouse diet 9F 5020; PMI Nutrition International), and water ad libitum. The Lox-Stop-Lox (*LSL*) *Kras^G12D/+^* (B6.129S4-*Kras* tm4Tyj/J #008179), and *Pdx1-Cre* (B6.FVB-Tg(*Pdx1*-cre)6Tuv/J) mice were purchased from the Jackson Laboratory (Bar Harbor, ME) and were bred to develop *LSL-Kras^G12D/+^;Pdx1-Cre* (KC) mice. The *B6.Cg-Gt(ROSA)26Sor^im14(CAG-tdTomato)Hze^/J* or Ai14 mice (#007914) were purchased from the Jackson Laboratory. The Ai14, *Pdx1-Cre* and *LSL-Kras^G12D/+^* were bred to develop *Rosa26^LSL-tdTomato^;LSL-Kras^G12D/+^; Pdx1-Cre* (AiKC) mice. CD45.1 (B6.SJL-Ptprca Pepcb/BoyJ) mice were purchased from the Jackson Laboratory (Bar Harbor, ME). Genotyping was performed according to Jackson Laboratory’s protocols (*Cre:* Protocol #21298, *Kras:* Protocol #29388 and Ai14: Protocol #29436). Activated *Kras* genotyping was performed as previously published.^1,3^ All mice were housed under identical conditions and are congenic on a C57BL/6J background (backcrossing > 15 generations). Both male and female mice of all strains were used. Original observations to develop the 9-month survival rate was based on 47 KC and 30 control genotypes at 9 months of age. After determining the etiology of death in the KC mice, there were 5 KC mice (out of 25 KC mice) that were identified as deceased from leukemia. We used 6 AiKC mice to visually confirm the location of *Pdx1-Cre* (projected location of mutant *Kras* expression) in the thymus, spleen and bone marrow at 12 weeks of age. We used 20 CD45.1, 3 AiKC mice and 3 C57BL6/J (CD45.2) mice at age 8 weeks for the bone-marrow transplant studies. The health and well-being of the mice were monitored closely by research and veterinary staff. Mice that showed signs of distress were immediately euthanized through CO^2^ asphyxiation.

### Histology

Thymus, pancreas, liver, spleen, and bone marrow from mice were collected and formalin-fixed and paraffin embedded (FFPE) by standard methods. Thymus samples were sectioned at 4 μm thickness and 50 μm intervals for a total of 6 slides. Pancreas, liver, spleen, and bone marrow were sectioned at 4 μm thickness and 200 μm intervals for a total of 10 slides. Sectioning and hematoxylin and eosin (H&E) staining was performed by the University of Wisconsin Experimental Animal Pathology Lab (EAPL) core facility. All histology was evaluated by a board certified gastrointestinal pathologist (KAM) or hematopathologist (EAR) as appropriate.

### CD3 Immunohistochemistry

CD3 staining was performed using the automated immunohistochemistry machine, the Ventana Discovery Ultra BioMarker Platform. Deparaffinization and heat-induced epitope retrieval in the form of “cell conditioning” with CC1 buffer (Ventana #950-500), a Tris based buffer, for approximately 32 minutes at 95^°^C was performed on the machine. Primary antibody anti-CD3 (2GV6) (Ventana #790-4341) was added undiluted and incubated for 20 minutes at 36^°^C followed by rinse with Reaction Buffer (Ventana #950-300). The secondary antibody, Discovery Omni-Map anti-Rabbit HRP (Ventana #760-4311) was added and incubated for 16 minutes at 37^°^C followed by rinse with Reaction Buffer (Ventana #950-300). Finally, the Discovery ChromoMap DAB detection kit (Ventana #760-159) was applied according to manufacturer instructions. The samples were removed from the instrument and rinsed with dawn dish soap, warm tap water and dH2O. Lastly, the harris hematoxylin counterstain 1:5 was applied for 45 seconds, followed by dH2O rinse, dehydration to xylene and cover slip.

### Genotyping for the activation of *Kras^G12D^* mutation construct

Activated *Kras^G12D^* refers to the successful Cre-mediated excision of the *Lox-Stop-Lox* sequence, allowing for transcription of the mutant *Kras* allele. To determine the penetration of *Kras^G12D^* mutation in thymuses of WT, KC and bone marrow chimeric mice, we followed the genotyping method for *Kras^G12D^* mutation previously described.^3^ Genomic DNA was isolated from formalin fixed paraffin embedded thymus slides using ReliaPrep FFPE gDNA MiniPrep System (Promega, Madison, WI). The DNA was then amplified using polymerase chain reaction (PCR) with the following probes: 5’-GGGTAGGTGTTGGGATAGCTG-3’ (OL8403) and 5’-CCGAATTCAGTGACTACAGATGTACAGAG-3’ (OL8404) with conditions previously published.^3^ These primers amplified a 325 bp band corresponding to the activated *Kras^G12D^* mutant allele and a 285 bp band corresponding to the WT allele.

### tdTomato immunohistochemistry (IHC)

AiKC mouse liver, thymus and spleen samples were formalin-fixed and paraffin embedded. Sectioning was performed by the University of Wisconsin Experimental Animal Pathology Lab (EAPL) core facility. Immunohistochemical staining for red fluorescence protein (tdTomato) was also performed by the EAPL. Sections were deparaffinized in xylenes and hydrated through graded alcohols to distilled water. Antigens were retrieved using citrate buffer pH 6.0 (10 mM Citric Acid, 0.05% Tween 20). Endogenous peroxidase was blocked with 0.3% H_2_O_2_ in PBS for 10 minutes at room temperature and blocking of non-specific binding was performed using 10% goat serum. Sections were incubated with rabbit anti-RFP antibody (600-401-379, Rockland Inc, Pottstown, PA) (1:1600) overnight at 4°C. After rinsing, sections were incubated with ImmPRESS goat anti-rabbit IgG (MP-7451, Vector Laboratories, Burlingame, CA) for 30 minutes at room temperature. Detection was performed using DAB substrate kit (8059S, Cell Signaling Technology, Danvers, MA). Samples were counterstained using Mayer’s hematoxylin (MHS32, Millipore-Sigma, St. Louis, MO) for one minute.

### tdTomato Fluorescent Visualization

AiKC mouse bone marrow was formalin-fixed and frozen embedded. Sectioning was performed by the University of Wisconsin EAPL core facility. Nuclei were stained with 4’, 6-diamidino-2-phenylindole (DAPI). Confocal images were acquired x10 objectives by a Zeiss Axio observer fluorescence microscope. Excitation was achieved using a Zeiss Colibri 7 (DAPI: 385 nm, tdTomato: 555 nm). The emitted fluorescence was collected with emission filters of DAPI (370-400 nm) and AF546 (540-570 nm).

### RNA isolation

RNA isolation was performed as previously described.^3^ Briefly, samples were stored in RNAlater (ThermoFisher Scientific, Waltham, MA) until RNA isolation (24 hours later). Following Qiazol lysis, samples were homogenized using a tissue homogenizer (Brinkmann Instruments, Model PT 10/35, 110 Volts, 6 Amps, 60 Hz) for spleen and thymus while RLT (Qiagen, Hilden, Germany) and hand homogenization was used to isolate BM. RNA was isolated using the Qiagen RNeasy Kit (Qiagen, Hilden, Germany). The extracted RNA was quantified using a spectrophotometer (ClarioStar Plate Reader, BMG LABTECH, Ortenberg Germany) and diluted to 50 ng/uL.

### Quantitative reverse transcription PCR

The qPCR was done as previously described.^3^ Briefly, 500 ng of RNA was reverse transcribed using the High Capacity cDNA Reverse Transcription Kit (Thermo Fisher, Waltham, MA) per manufacturer protocol. The qPCR was performed on the Thermo Fisher QuantStudio 7 (Thermo Fisher, Waltham, MA). All reactions were run in triplicate. Results were analyzed using the delta-delta CT method. The following TaqMan^®^; probes were used: *Cre* (Enterobacteria phage P1 cyclization recombinase, Mr00635245_cn), *Pdx1* (pancreatic and duodenal homeobox 1, Mm00565835_cn) and the house keeping gene Hprt (hypoxanthine guanine phosphoribosyl transferase, Mm03024075_m1) (Thermo Fisher, Waltham, MA).

### Flow Cytometry

Bone marrow was collected from the femurs and tibias of AiKC mice and filtered through a 70 μm cell strainer. The single cell suspension was adjusted to 1 million cells per sample, suspended in 400ul of staining buffer (IMDM + 10% FBS) and stained with an antibody cocktail (supplemental table 1). Stained cells were run using the Attune Nxt (ThermoFischer) and analyzed using FlowJo. The following gating strategy was used to delineate live cells: FSC-A X SSC-A to FSC-A X FSC-H to SSC-A X SSC-H to FSC-A X DAPI. Live cells are DAPI negative.

To identify bone marrow subsets from the live cells the following gating strategy was used (supplemental figure 1): FITC x FSC (FITC negative = lineage negative cells) to Kit x Sca1 (gated on c-Kit^+^Sca1^+^) to CD150 negative (multipoint progenitor cells) and CD150^+^ (hematopoietic stem cells). All final subsets were analyzed as FSC-A x tdTomato to identify the tdTomato positive cells in each group. Statistical comparisons were made using Fisher’s Exact test (significance was assigned for P-values < 0.05).

To determine the donor chimerism, peripheral blood was collected in blood collection tube with K_2_EDTA (BD Microtainer tubes), and the red blood cells were lysed with RBC lysis buffer (Santa Cruz Biochemicals, Dallas, TX). The leukocytes were labelled with CD45.1-PE (Biolegend, 12-0453-81) and CD45.2-FICT (Biolegend, 11-04540-81) antibodies. The cells were analyzed using the Attune Nxt (ThermoFischer) and analyzed using FlowJo.

### Bone Marrow Transplant

B6 CD45.1 recipient mice were sub-lethally irradiated (5.5 Gy, twice) at 9-10 weeks of age. Bone marrow cells from 3 pooled donor mice (C57BL6/J mice or AiKC mice) were isolated and bone marrow transplantation performed via retro-orbital sinus injection. For instance, 1 × 10^7^ bone marrow cells pooled from three separate AiKC mice were transplanted (injected via retro-orbital sinus) into ten B6 CD45.1 mice. Following transplantation, the recipient mice (immunosuppressed due to sub-lethal irradiation) were placed on a Uniprim diet containing 275 parts-per-million (ppm) trimethoprim and 1365 ppm sulfadiazine (Test Diet Company TD.06596) for 3 weeks post-irradiation in place of standard chow. Recipient mice were monitored closely for 3 weeks following transplantation to ensure no lethality related to the transplant procedure. Retro-orbital bleeds were performed to assess for donor chimerism. Mice were evaluated until time of onset of leukemia as evidenced by signs of duress or respiratory distress, or until the time of study end-point at 5 months post-transplant.

## RESULTS

### Development of T-ALL in the KC mouse model

The KC mouse model is generated by crossing the LSL-*Kras^G12D/+^* (K) mouse to pancreatic and duodenal homeobox 1 (*Pdx1)-Cre* (C) mouse and it is typically used to evaluate pre-neoplastic pancreas lesions where *Kras*-mutation is targeted to pancreas cells. Consistent with current literature, the KC mice evaluated in this study developed the expected rates of pancreatic cancer precursor lesions called pancreatic intraepithelial neoplasia (PanIN), with 100% (36/36) of KC mice developing PanIN-1, 5.6% (2/36) developing PanIN-2, and 8.3% (3/36) developing PanIN-3. Moreover, as expected, approximately 13.9% (5/36) developed pancreas cancer and 36.1% (13/36) developed surface tumors called papillomas. Original observations of survival out to age 9 months revealed a 70.2% (33/47) survival rate, with eleven of the KC mice unable to be evaluated due to being found deceased (age range 14.1 – 29.1 weeks). Meanwhile, three mice were identified at moribund status (>20% weight loss and slow movements) and were euthanized (age 26.7 weeks, 16 weeks and 15.7 weeks). Two of the moribund mice had abdominal cavity evaluated at necropsy and no abnormalities were identified; the pancreas demonstrated only chronic pancreatitis and PanIN-1 with mostly preserved pancreatic architecture. Thus, it was unclear at the time the cause of moribund status. Subsequently, the third moribund KC mouse (age 15.7 weeks) that was identified demonstrated chest bulging at necropsy and thus the thoracic cavity was evaluated. This revealed a large thymic tumor (abdominal exploration revealed no abnormalities and pancreas demonstrated chronic pancreatitis / PanIN-1). Subsequently, mice were closely evaluated for either weight loss or respiratory changes and an additional 5 (out of 25) KC mice were found to develop respiratory distress and/or weight loss prompting euthanization which revealed large thymic tumors on necropsy (figure 1A, B). On histopathologic examination of the thymic tumors, there was a uniform infiltration of immature blast cells with frequent mitotic figures, consistent with T-ALL (figure 1C). Finally, the diagnosis of T-ALL was confirmed by expression of cytoplasmic CD3 (figure 1D).^4^ The pancreas, stomach, small intestine, colon, spleen, thymus, lungs and liver underwent gross analysis, but no other solid organ manifestation of T-ALL was readily apparent in the KC mice. Nor was there any evidence of pancreas pathology (beyond PanIN-1) that could have precipitated duress in these mice. Only KC mice developed tumors (i.e. only mice possessing activated *Kras*-mutation), while no age-matched *LSL-Kras^G12D/+^* (n = 10), *Pdx1-Cre* (n = 10) or C57BL/6J (n=10) mice developed leukemia, Pan-IN, pancreas cancer or papillomas, and 100% of control mice lived to age 9 months.

**Figure 1:**
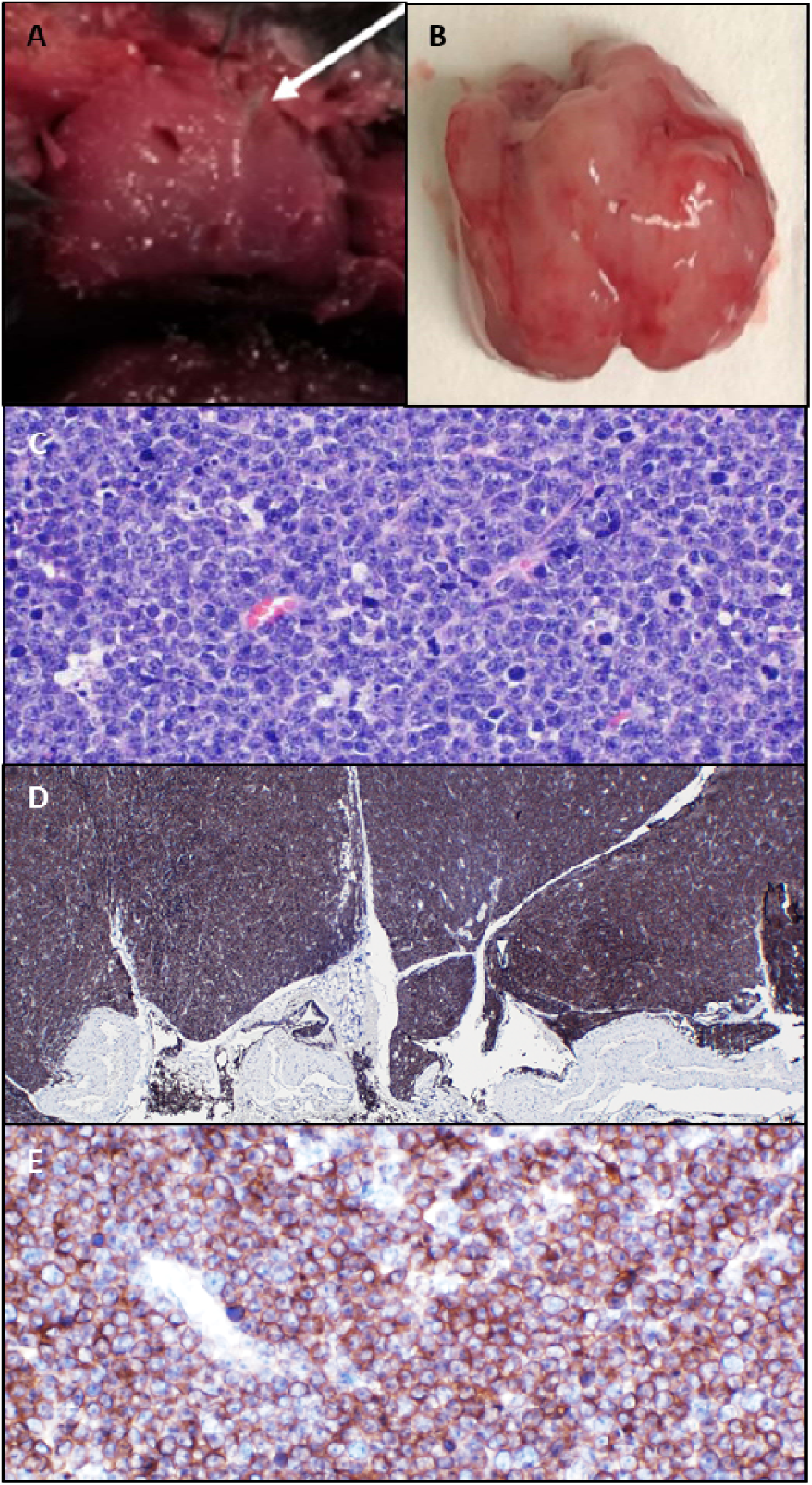
T-ALL driven by activated *Kras*-mutation. KC mice develop thymic masses, demonstrated *in situ* (A), and *ex-vivo* (B). On histopathologic analysis (H&E), there is diffuse infiltration of immature blast cells with convoluted nuclear membrane, indistinct nucleoli and frequent mitotic figures (C, 400x magnification). These abnormal appearing cells obliterate the normal cortex and medulla and are positive for cytoplasmic CD3, confirming T-ALL diagnosis (D, 40x magnification; E, 400x magnification).

### T-ALL development in KC mouse model is driven by mutated-*Kras* gene

*Kras* is the most common mutation in cancer and expression of *Kras* in other tissues can drive cancer formation. For instance, we identified a sex-dependent development of *Kras*-driven anal SCC in KC mice.^3^ Given that the thymic tumors only developed in mice harboring a conditional mutant-*Kras^G12D^* gene, we hypothesized that the *LSL-Kras^G12D/+^* gene was being activated in the hematopoietic cells driving the development of T-ALL (and the concomitant thymic tumor). To detect the activated *Kras^G12D^*-mutation (*L-Kras^G12D^)*, DNA was isolated from KC normal thymus and KC thymic tumor and was assessed for activated *Kras*-mutation (i.e. *Cre*-mediated excision of *Lox-Stop* sequence). KC mice that did not manifest thymic tumor (i.e. normal thymus) were used as a ‘negative control’ because these mice possessed the mutant *Kras* allele and *Pdx1-Cre* transgene, but did not demonstrate evidence of leukemia. PCR analysis of the genomic DNA demonstrated *L-Kras^G12D^* in the thymic tumors of KC mice, but only wild type *Kras (Kras^+^*) in normal thymus of KC mice without T-ALL (Figure 2). This finding indicates the T-ALL / thymic tumors are driven by the mutated *Kras^G12D^* gene.

**Figure 2:**
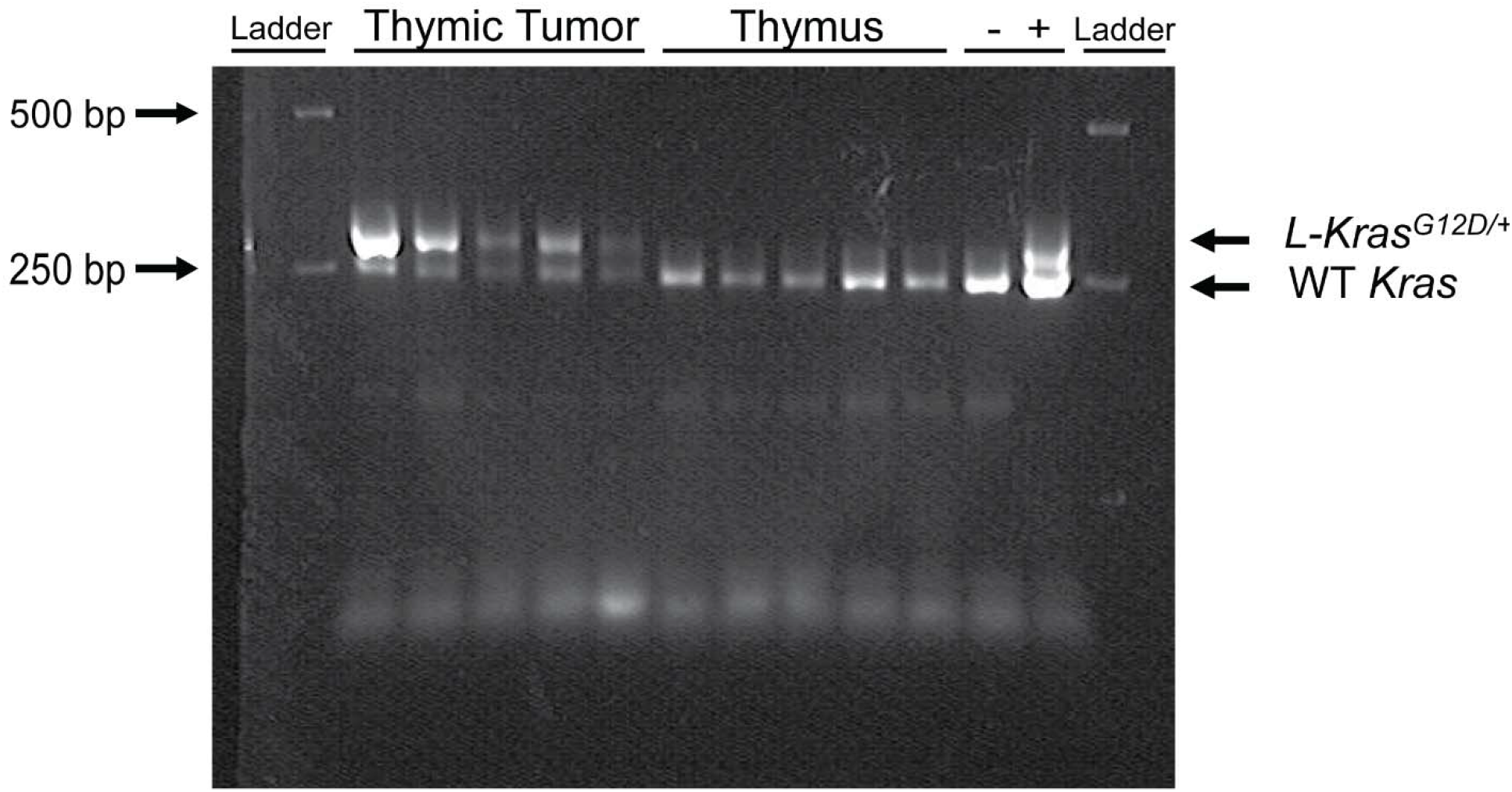
The activated *Kras*-mutation is present in thymic tumors but not normal thymus. PCR of genomic DNA from thymus of KC mice without T-ALL (Thymus) reveals wild type *Kras* (WT *Kras)*. In KC mice with T-ALL, the active form of *Kras*-mutant allele (*L-Kras^G12D/+^*) is present in the thymic tumors (tumor), indicating *Cre*-mediated excision of Lox-Stop sequence. Positive control (+) consisted of splenocytes from *Vav-Cre; LSL-Kras^G12D/+^* mice (as previously described)^3^ and negative control (-) was tail from C57BL/6J mice. These are shown along with a ladder demonstrating the 250 and 500 bp markers.

### *Pdx1-Cre* is expressed in the thymus, spleen and bone marrow

The *Kras^G12D^* mutation is activated by *Pdx1* driven *Cre* indicating *Pdx1* expression outside of the pancreas causes leukemia development in KC mice. While *Pdx1* has traditionally been regarded as pancreas-lineage specific, evidence clearly demonstrates *Pdx1* expression in other organ systems. For instance, we identified *Pdx1* expression in anal epithelium,^3^ and evaluation of human and mouse hematologic cell databases reveals *Pdx1* expression in hematopoietic cells.^5, 6^ To ascertain where *Pdx1-Cre* is being expressed within the KC mouse model, we crossed the KC mice to Ai14 mice to generate the AiKC ‘marker’ mouse model (*Rosa26^LSL-tdTomato^; LSL-Kras^G12D/+^; Pdx1-Cre)*. AiKC mice harbor the *LSL-tdTomato* red fluorescent gene in the *Rosa26* locus, and in the presence of *Cre*-recombinase, the stop sequence is excised allowing for expression of tdTomato protein, localizing *Pdx1 (Pdx1-Cre*) expression and functioning as a marker for expression of mutant-*Kras^G12D^* gene. Through IHC staining for tdTomato, we found *Pdx1-Cre* was expressed in pancreas as expected, and also in thymus and spleen (Figure 3 A,B,C,D,E,F). In conjunction, immunofluorescence (IF) demonstrated *Pdx1* expression in thymus, spleen, liver and high expression in bone marrow, which contains the hematopoietic cells (Figure 3 G,H,I). This was further confirmed through qPCR showing *Pdx1* expression in these tissues of WT and KC mice (Figure 3 J, K).

**Figure 3:**
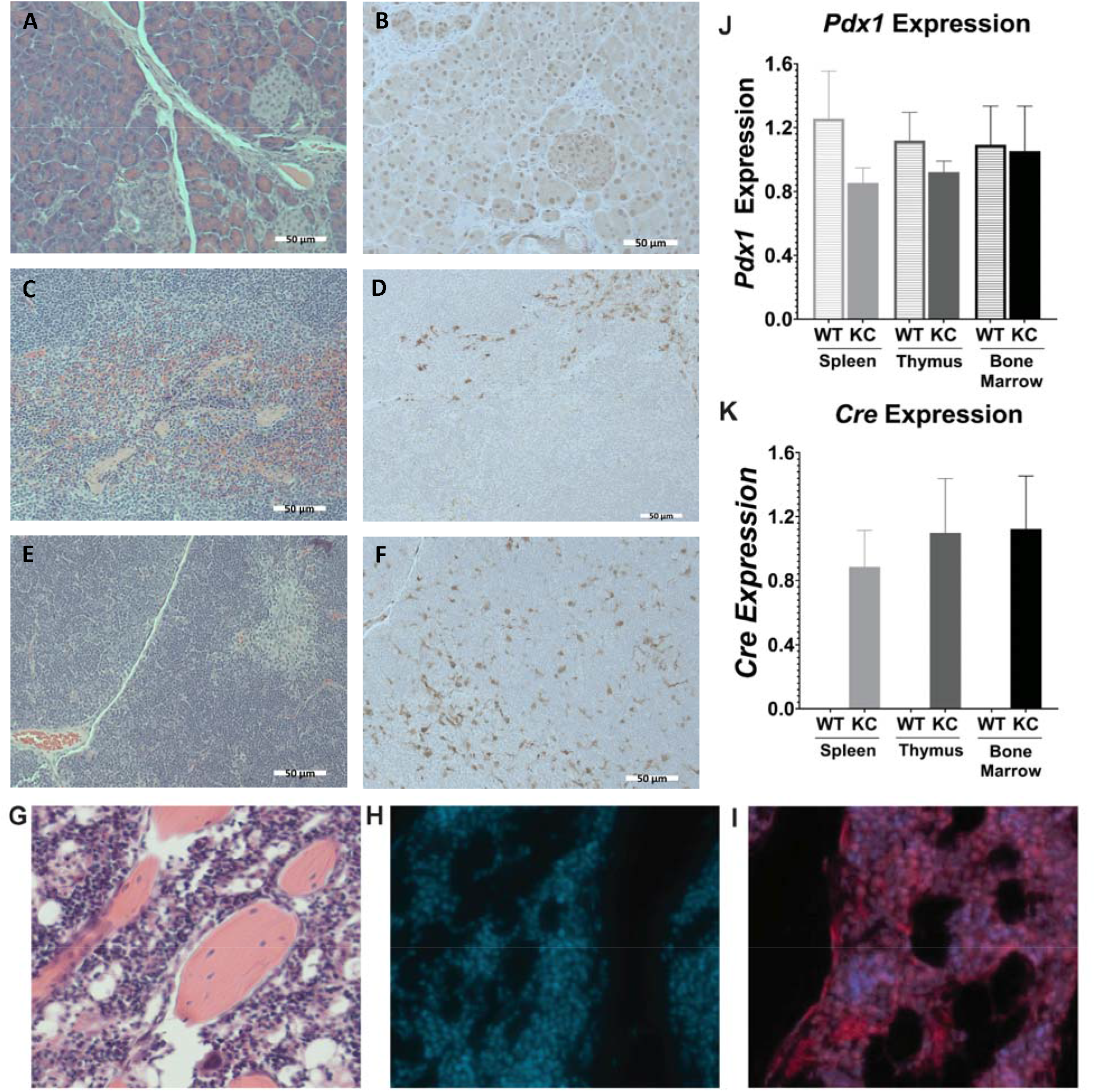
*Pdx1-cre* expression in pancreas, spleen, thymus and bone marrow. AiKC mice *LSL-Tdtomato* (*Rosa26*;KC) express tdTomato in cells expressing *Pdx1*. Pancreas (A,B), spleen (C,D), thymus (E,F), and femur (G,H,I) H&E (A,C,E,G) with anti-tdTomato antibody (B,D,F brown dots) demonstrate cells with *Pdx1* expression. Immunofluorescence shows DAPI stained cells (H) and DAPI with tdTomato (I), confirming *Pdx1* expression in the bone marrow. *Pdx1* (J) and *Cre* (K) expression was confirmed using qPCR. Scale bars equal 50 μm.

### *Pdx1* is expressed in multipotential progenitor cells (MMP)

The prevalence of *Pdx1* expression in immune organs (in particular the bone marrow) led us to determine which cell types in the bone marrow preferentially express *Pdx1*. Bone marrow was isolated from AiKC mice and analyzed for *Pdx1* expression using flow cytometry including the lineage-committed cells, the lineage negative cells, the multipotential progenitor cells (MPP) and hemopoietic stem cells (HSC). The lineage committed cells consist of bone marrow cells that have committed to develop into a specific mature immune cell. The lineage negative cells encompass cells that have not committed to their immune lineage and are either MPP or HSC. MPPs are able to differentiate into any immune cell type provided the right stimulus is present, but cannot self-renew. Finally, the HSCs are stem cells that can either undergo self-renewal (i.e. generate more HSCs) or differentiate into progenitor immune cells and blood cells. HSCs are also characterized by their ability to repopulate the immune cell compartment and are therefore considered bone marrow ‘repopulating’ cells.^7^ We found that *Pdx1* was statistically significantly expressed in the bone marrow cells, lineage negative cells, and MPP cells and not within the lineage positive cells or HSC population (Figure 4). This indicates that *Pdx1* allows for expression of *Cre* which activates the mutated-*Kras^G12D^* gene in these cells, thus likely driving the development of the T-ALL found in these mice. This finding of *Pdx1* expression in KC mice aligns with the known expression of *Pdx1* in murine hematopoietic lineage cells such as common lymphoid progenitors and T-cells.^5^ The expression levels seen in murine hematologic cells is also concordant with human expression of *Pdx1* in hematologic cells.^5, 6^

**Figure 4:**
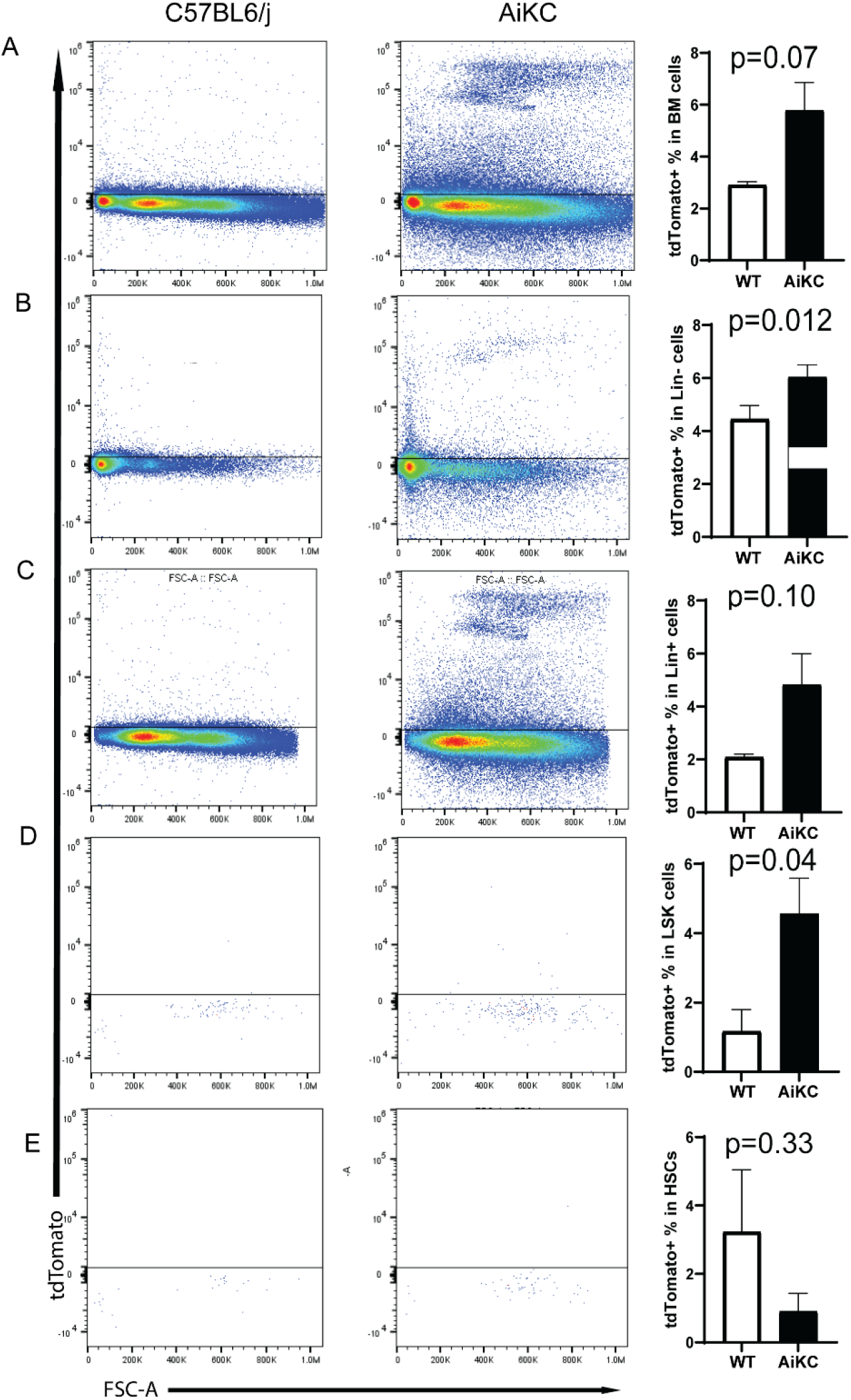
*Pdx1* expression within the multipotential progenitor cells in the bone marrow. Bone morrow was isolated from the AiKC mice and the cells expressing *Pdx1* were analyzed. (A) The expression level of tdTomato cells in the bone marrow (BM) (P=0.07). (B)There is significant expression of tdTomato cells in lineage negative BM cells (P=0.012), (C) but not lineage positive BM cells (P=0.10). (D)There is significant expression in the multipotential progenitor cells (P=0.04). (E) There is not significant expression within the hematopoietic stem cells (P=0.33).

### Bone marrow transplant confirms hematopoietic etiology of T-ALL

*Kras*-mutant leukemia developing in KC mice is driven by *Pdx1* expression and Cre mediated excision of lox-stop, and we hypothesized that the hematopoietic compartment (as opposed to spleen or thymus) likely contains the cell of origin for the T-cell malignancy seen in these mice. To confirm, we performed a bone marrow transplant (BMT) study where pooled bone marrow collected from three AiKC mice was transplanted into irradiated B6 CD45.1 mice (AiKC mice express CD45.2). The AiKC mice were otherwise healthy 8-week-old mice with no evidence of pathology on gross analysis. As a control, pooled bone marrow from three B6 CD45.2 mice was transplanted into irradiated B6 CD45.1 mice. No mice in either group died as a result of toxicity related to the irradiation or BMT. We found 100% (10/10) of the CD45.1 mice transplanted with AiKC bone marrow died of their disease by 4 months post-transplant (Figure 5). All deceased mice demonstrated large thymic tumors with uniform cellular infiltration and presence of *Kras* mutation confirmed by PCR analysis (Figure 6). In contrast, none (0/8) of the age matched CD45.1 mice transplanted with C57BL6/J (CD45.2) bone marrow died. Flow cytometry confirmed successful transplant of the CD45.2 bone marrow into CD45.1 mice (Supplemental Figure 2), and thymus from these wild-type BMT controls demonstrated normal histopathology. These results confirm that *Pdx1* expression in the hematopoietic compartment promotes Cre expression (*Pdx1-Cre* transgene) in KC mice, resulting in activation of mutant *Kras^G12D^* in hematopoietic cells and development of T-ALL, with subsequent death in the afflicted KC mice.

**Figure 5:**
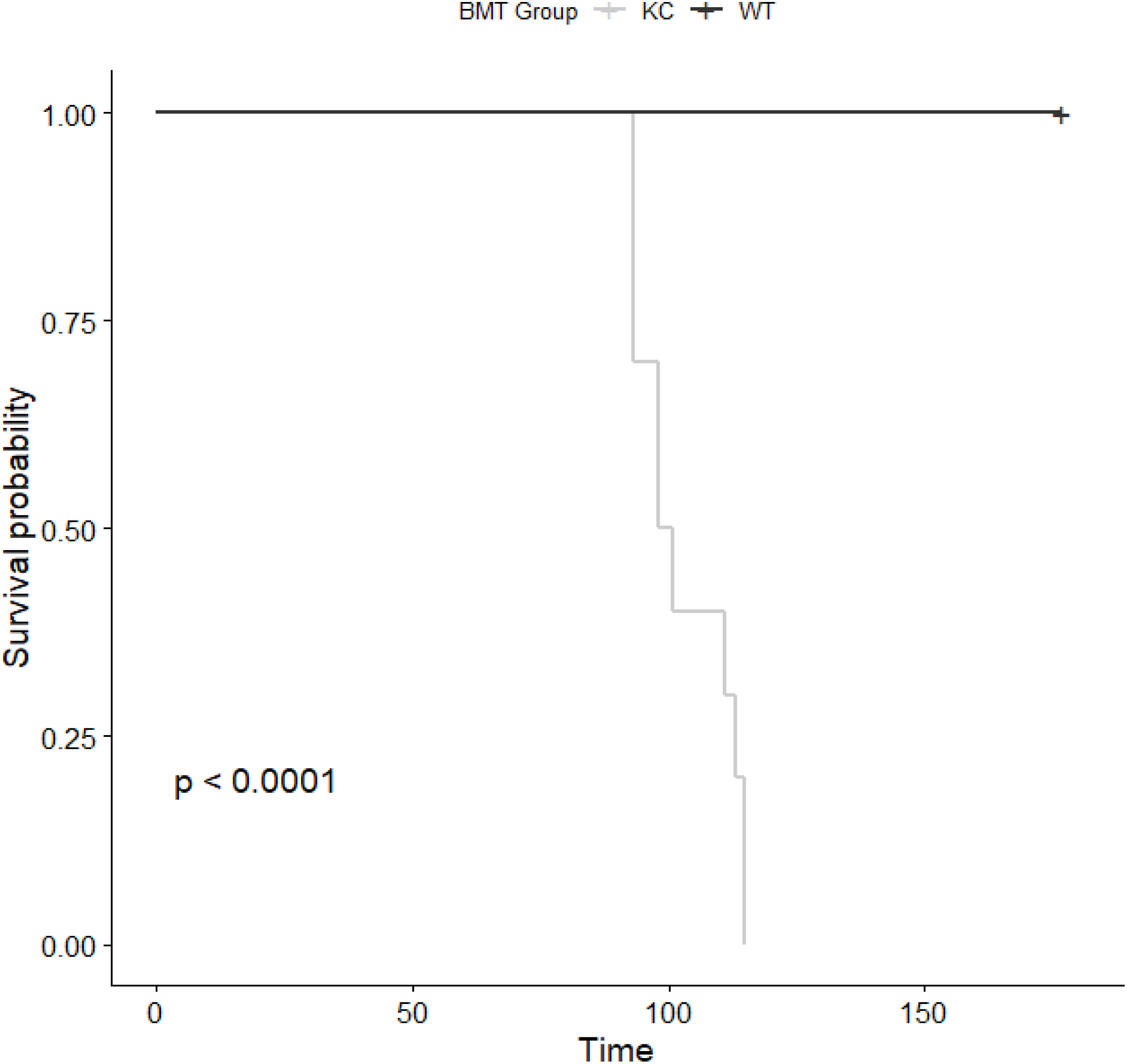
Kaplan-Meier survival curve in bone marrow transplant mice. Bone morrow was isolated from three separate AiKC mice and pooled for transplant via retro-orbital injection into irradiated CD45.1 B6 mice. Additionally, bone marrow was isolated from three separate C57BL/6J (CD45.2) mice and pooled for transplant into irradiated CD45.1 B6 mice. Following transplant, mice were monitored for signs of duress. There were no deaths in the immediate post-transplant period. Time of survival following BMT was measured and recorded. Onehundred percent of the CD45.1 mice receiving AiKC bone marrow (KC: light grey) died due to TALL and thymic tumor development within 4 months of transplant, compared to none of the CD45.1 mice receiving wild-type bone marrow (WT: dark grey) [p<0.0001].

**Figure 6:**
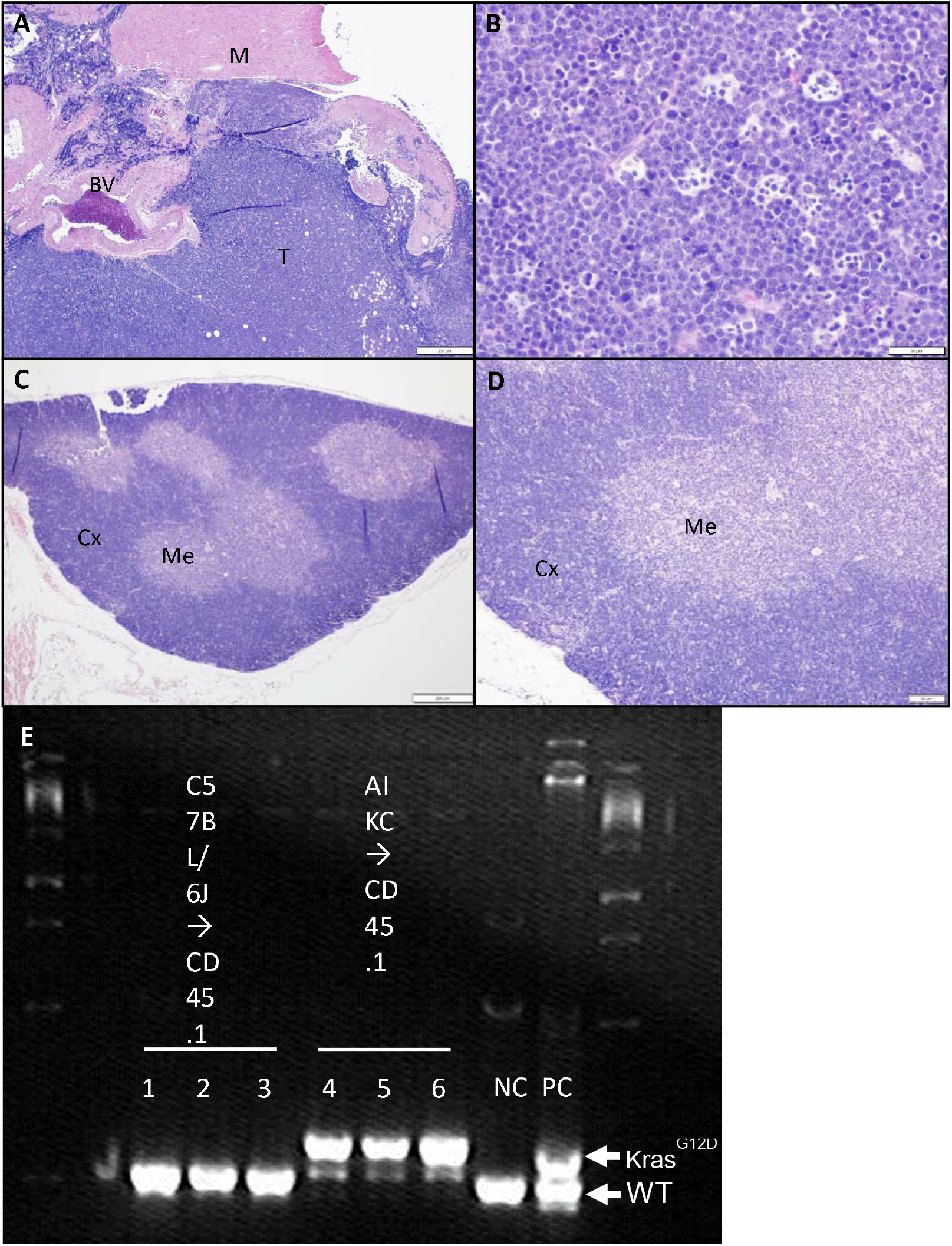
Transplant of KC bone marrow into wild-type results in *Kras*-mutant T-ALL. Pooled bone marrow from three KC mice was transplanted into sub-lethally irradiated B6 CD45.1 mice. Thymic tumors in recipient mice were removed for histopathologic analysis. Demonstrated is the thymic tumor (T) adjacent to major blood vessels (BV) and myocardium (M) [40x; scale bar = 200 μm]. On closer magnification (B), there is diffuse infiltration of immature T-cells with convoluted nuclear membrane, indistinct nucleoli and frequent mitotic figures, consistent with T-ALL seen primarily in the KC mice (200x; scale bar = 20 μm). Histopathology from CD45.1 mice transplanted with C57BL6/J bone marrow revealed normal thymus containing the cortex (Cx) and medulla (Me) [40x; scale bar = 200 μm] (C). These epithelial compartments are also evident on higher magnification (D) [100x; scale bar = 50 μm]. DNA isolated from thymus of CD45.1 recipients of C57BL6/J bone marrow (1,2, 3), and KC bone marrow (4, 5, 6) and demonstrated wild-type *Kras* and activated *Kras*-mutation, respectively (E). Positive control (PC) consisted of splenocytes from *Vav-Cre; LSL-Kras^G12D/+^*mice (as previously described)^3^ and negative control (NC) was tail from C57BL/6J mice.

## DISCUSSION

The KC pancreas-cancer pre-cursor mouse model (i.e. PanINs) was first developed in 2003^1^ and has since continued to be highly utilized in pancreas cancer studies. Often, the *Pdx1-Cre* driven *Kras^G12D^*-mutant (KC) mouse is typically crossed to mice harboring additional genetic mutations (e.g., Trp53, Ink4a/Arf) or mice are subjected to carcinogenic environmental changes (e.g., high-fat diet) to explore advanced pancreas cancer.^8–10^ Throughout the use of this mouse model, the published survival of roughly 70% of KC mice at 9 months has been attributed to pancreas cancer.^2^ However, in KC mice, post-mortem evaluation of the pancreas proves difficult due to organ auto-digestion. Notably, while we did not identify evidence for metastatic pancreas cancer in moribund mice or at necropsy, we did find mice have evidence of advanced hematologic cancer (T-ALL) resulting in a large thymic tumor which caused respiratory suppression as the cause of death. This novel finding is extremely important to consider when using this mouse model in pancreas cancer studies at advanced age (i.e. 9 months age), especially when planning to evaluate immunologic contributions to pancreas cancer development. Under these circumstances, there could be a competing cause for lethality (i.e. exacerbation of leukemia phenotype), or immunologic alterations that may not be attributable to pancreas cancer development but instead due to mutant-*Kras* driven changes in the hematopoietic compartment.

We used pooled bone marrow from 3 AiKC mice to transplant into 10 mice because roughly 30% of mice were deceased of leukemia by 9 months in the KC mice, so we suspected pooling of three separate samples would heighten the likelihood of hematopoietic cells containing mutant-*Kras*. Thus, this pooling of bone marrow resulted in a ‘solitary sample’ with activated *Kras^G12D^*, and explains the difference between the BMT mice (where 100% developed T-ALL) and the KC mice (where 30% die by age 9 months). Similar to other *Kras*-mutant leukemia models,^11, 12^ BMT mice receiving AiKC bone marrow were deceased within 3-4 months of transplantation. This reflects the aggressiveness of *Kras*-mutant leukemia, where onset of *Kras*-mutation in the hematopoietic cells causes rapid demise. In fact, in the KC mice that were found to harbor thymic tumors with T-ALL, the age of onset was roughly 14-29 weeks, with a median age of 18 weeks. Moreover, in KC mice that were spontaneously deceased in this study (prior to identification of the leukemia phenotype), all were found deceased near the median age of 18 weeks. When evaluating the pancreas of mice at age 4.5 months,^1^ the majority of the pancreas is normal with few foci of the earliest pancreatic pre-cancerous lesion (PanIN-1), thus further refuting an underlying pancreas pathology as the cause of early death in KC mice.

The aggressive nature of the *Kras*-mutant leukemia manifest in the KC mice (deceased within 3-4 months of BMT) is also reflected in human disease. T-cell acute lymphoblastic leukemia is a heterogeneous and lethal malignancy of immune cells that accounts for 10-15% of pediatric ALL, and 25% of adult ALL. Although high intensity combination chemotherapeutic regimens have success in pediatric and adult cases, ~25% of pediatric and ~50% of adult patients fail initial therapy or relapse.^4^, ^13^ Prognosis is poor in these patients, with 80% mortality in spite of aggressive and toxic salvage protocols (e.g. intensive chemotherapy followed by allogeneic stem cell transplant).^14^ These dismal outcomes reflect the highly resistant nature of refractory or relapsed disease, with only 30-40% of patients responding to second line therapy, yielding a median survival of 6 months.^13^ Notably, *Kras*-mutation is disproportionately present in recurrent disease (40%) in comparison to newly diagnosed T-ALL (5-10%),^15^ providing insight into refractory T-ALL. Thus a *Kras*-driven mouse model of T-ALL provides an excellent platform to understand high-risk T-ALL (i.e. refractory disease).^16^ Moreover, because there is not full penetrance with the KC model (not all KC mice developed overt hematologic malignancy), this model could be considered to evaluate factors (e.g. transcription factors) that exacerbate development of *Kras*-mutant T-ALL. For example, a direct 1:1 BMT of KC mice into CD45.1 mice (i.e. not pooling of bone marrow as was done in this study) should result in only ~20-30% of mice developing leukemia, and allow evaluation of transcription factors that precipitate *Kras*-mutant leukemia development (while also circumventing the ‘competing’ pancreas pathology).

It is important to note that not all mice were able to be evaluated either at age 9 months, or when captured at moribund status (weight loss, poor movement or respiratory distress), even after the point of close scrutiny when hematologic malignancy had been discovered in the KC mouse model. Some mice spontaneously died without obvious signs that precipitated the need for euthanasia, and these mice had undergone autodigestion of organs prior to effective evaluation. We identified 5 KC mice with thymic tumors from 25 KC mice that were evaluated, but there were five additional mice during this time period that were spontaneously deceased (i.e. ten out of thirty mice died prior to age 9 months). It is plausible that other hematologic malignancies (e.g. acute myelogenous leukemia) may arise in the KC mice that does not result in respiratory distress / thymic tumors, but that may cause spontaneous death. Evidence to support this is the fact that Pdx1 is expressed in the MPP cells and lineage negative cells in KC mice, which may result in other types of leukemia. However, additional studies are required to specifically address this current limitation.

## CONCLUSIONS

This study identified the spontaneous development of T-ALL in the KC mouse model, a model highly utilized in pancreatic cancer research. The development of T-ALL in the KC model stems from the expression of Pdx1 within the MPP cells of the BM driving the expression of the mutated *Kras^G12D^* gene in this cell population resulting in the development of T-cell leukemia. TALL, particularly refractory T-ALL, is a highly lethal cancer with 80% of those who relapse or fail first line therapies succumbing to their disease. Notably, *Kras*-mutation is disproportionately present in recurrent disease (40%) in comparison to newly diagnosed T-ALL (5-10%) presenting a unique etiology of the recurrent disease.^14^ Given the KC mice develop Kras driven T-ALL that is not 100% penetrant and is transplantable, this model has the potential to be utilized to investigate the onset and exacerbation of the T-ALL. Finally, the results of this study are highly important to those utilizing the KC model for pancreas cancer research and particularly for those investigating immunologic changes within this model.

## LIST OF ABBREVIATIONS

Ai14: B6.Cg-*Gt(ROSA)26Sor^tm14(CAG-tdTomato)Hze^*/J
AiKC: B6.Cg-*Gt(ROSA)26Sor^tm14(CAG-tdTomato)Hze^/J*; *Pdx1-Cre; LSL-Kras^G12D/+^*
BMT: Bone marrow transplant
FFPE: Formalin-fixed, paraffin embedded
H&E: Hematoxylin and eosin
HSC: Hematopoietic stem cells
IHC: Immunohistochemistry
KC: *Pdx1-Cre; LSL-Kras^G12D/+^*
LSL: Lox-stop-lox
MPP: Multipotent progenitor cells
PanIN: Pancreatic intraepithelial neoplasia
Pdx1: Pancreatic and duodenal homeobox 1
T-ALL: T-cell acute lymphoblastic leukemia

## DECLARATIONS

### Ethics approval and consent to participate

The use of animals for this study was approved by the IACUC

### Consent for publication

Not applicable

### Availability of data and materials

The data generated or analyzed during the study are included in the published article (and the supplementary information).

### Author’s contributions

MW contributed to the data acquisition (prepared Figure 1, Figure 3, Figure 5, supplemental table 1), data analysis and writing of the manuscript. MN contributed to the data acquisition (prepared Figure 2, Figure 3, Figure 4, Figure 6, supplemental figures 1 and 2), data analysis, interpretation of data, and editing of the manuscript. MW and MN contributed equally to this work as co-first authors. ER contributed to the data acquisition, data analysis and editing of the manuscript. KM contributed to the data acquisition, data analysis and editing of the manuscript. SRK designed the study, contributed to the data acquisition, data analysis, interpretation of data, and writing of manuscript.

## Acknowledgements

The authors would like to acknowledge the Experimental Animal Pathology Lab (EAPL) supported by the UWCCC (P30 CA014520) for use of its facilities and services.

