## Supplemental Figure 1 for "Pdx1 expression in hematopoietic cells activates *Kras*-mutation to drive leukemia in KC (*Pdx1-Cre ; LSL-Kras^G12D/+^*) mice"

**Supplemental Figure 1: Gating Strategy** After live cells are gated on, a gate is placed around the FITC negative population (A) to gate out the lineage positive cells. Then a gate is placed around the Kit+ and Sca1+ cells to delineate progenitor and stem cells (B). Finally, multipotential progenitor (MPPs) are CD150 – and the hematopoietic stem cells (HSC) are CD150+ (C). All groups are then plotted against tdTomato to identify the tdTomato expressing cells.

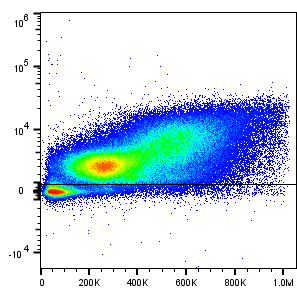

FSC-A

Lin CD48

Lin^-^

A)

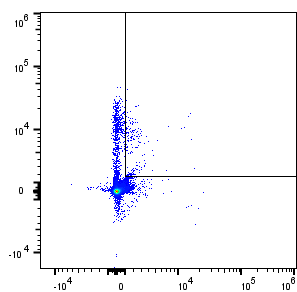

B)

Sca-1

c-Kit

Lin^-^Sca1^+^c-Kit^+^

(LSK)

C)

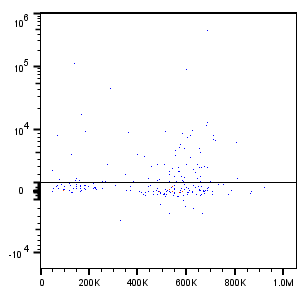

CD-150

FSC-A

MPP

HSC
