## Supplemental Figure 2 for "Pdx1 expression in hematopoietic cells activates *Kras*-mutation to drive leukemia in KC (*Pdx1-Cre ; LSL-Kras^G12D/+^*) mice"

**Supplemental Figure 2: Chimeric bone marrow in B6 CD45.1 recipients from C57BL6/J donors.** Flow cytometry of bone marrow from CD45.1 and C57BL6/J mice (A, B) demonstrate expected antigen detection. Pooled bone marrow from three C57BL6/J donors (CD45.2) were transplanted into sub-lethally irradiated B6 CD45.1 mice. The CD45.1 recipient mice were healthy with no evidence of pathology and were ultimately euthanized at 5-months post-BMT. Bone marrow from four of the mice was harvested and flow cytometry performed to confirm successful transplantation (C, D, E, F).

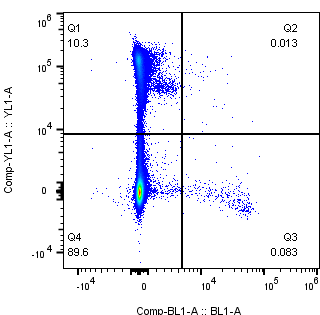

**CD45.1**

**CD45.2**

**CD45.1 Mouse**

**A**

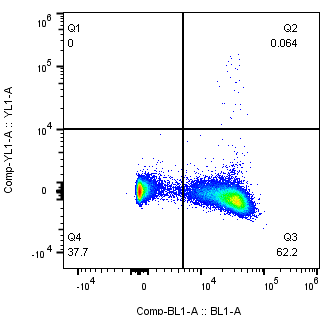

**CD45.1**

**CD45.2**

**C57BL/6 Mouse**

**B**

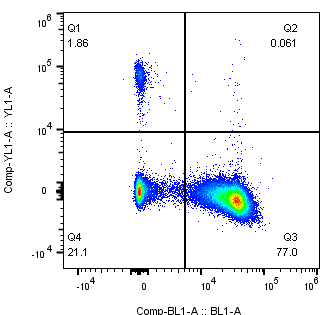

**CD45.1**

**CD45.2**

**C57BL6/J Donor- CD45.1 Recipient_1**

**C**

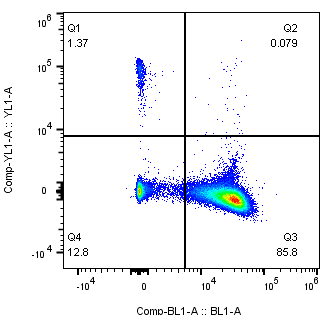

**CD45.1**

**CD45.2**

**C57BL6/J Donor- CD45.1 Recipient_2**

**D**

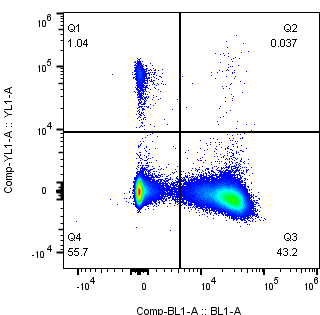

**C57BL6/J Donor- CD45.1 Recipient_3**

**CD45.1**

**CD45.2**

**E**

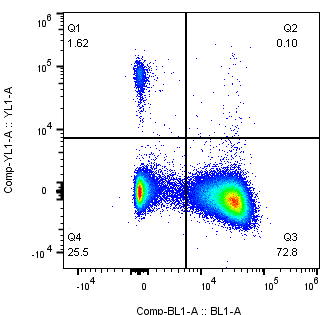

**CD45.1**

**CD45.2**

**C57BL6/J Donor- CD45.1 Recipient_4**

**F**
