## Supplemental Table 1 for "Pdx1 expression in hematopoietic cells activates *Kras*-mutation to drive leukemia in KC (*Pdx1-Cre ; LSL-Kras^G12D/+^*) mice"

| Supplemental Table 1: Flow cytometry antibody information | | | |
| --- | --- | --- | --- |
| Cell Marker | **Fluorochrome** | **Product information number** | **Dilution** |
| B220 | FITC | BD 11-0452 | 1:200 |
| CD3 | FITC | BD 11-0031 | 1:200 |
| CD4 | FITC | BD 11-0041 | 1:200 |
| CD5 | FITC | BD 11-0051 | 1:200 |
| CD8 | FITC | BD 11-0081 | 1:200 |
| CD48 | FITC | BD 11-0481 | 1:200 |
| Gr-1 | FITC | BD 11-5931 | 1:800 |
| TER-119 | FITC | BD 11-5921 | 1:100 |
| CD150 | PE-Cy7 | Biolegend 115914 | 1:200 |
| Sca-1 | PerCP-Cy5.5 | BD 45-5981 | 1:100 |
| c-Kit | APC | BD 17-1171 | 1:1000 |
| CD45.1 | PE | Biolegend, 12045381 | 1:100 |
| CD45.2 | FICT | Biolegend, 11-04540-81 | 1:100 |
| Live/Dead Stain | DAPI | ThermoFischer | 1:1000 |
